## Supplementary information for "Base-pair resolution analysis of the effect of supercoiling on DNA flexibility and major groove recognition by triplex-forming oligonucleotides"

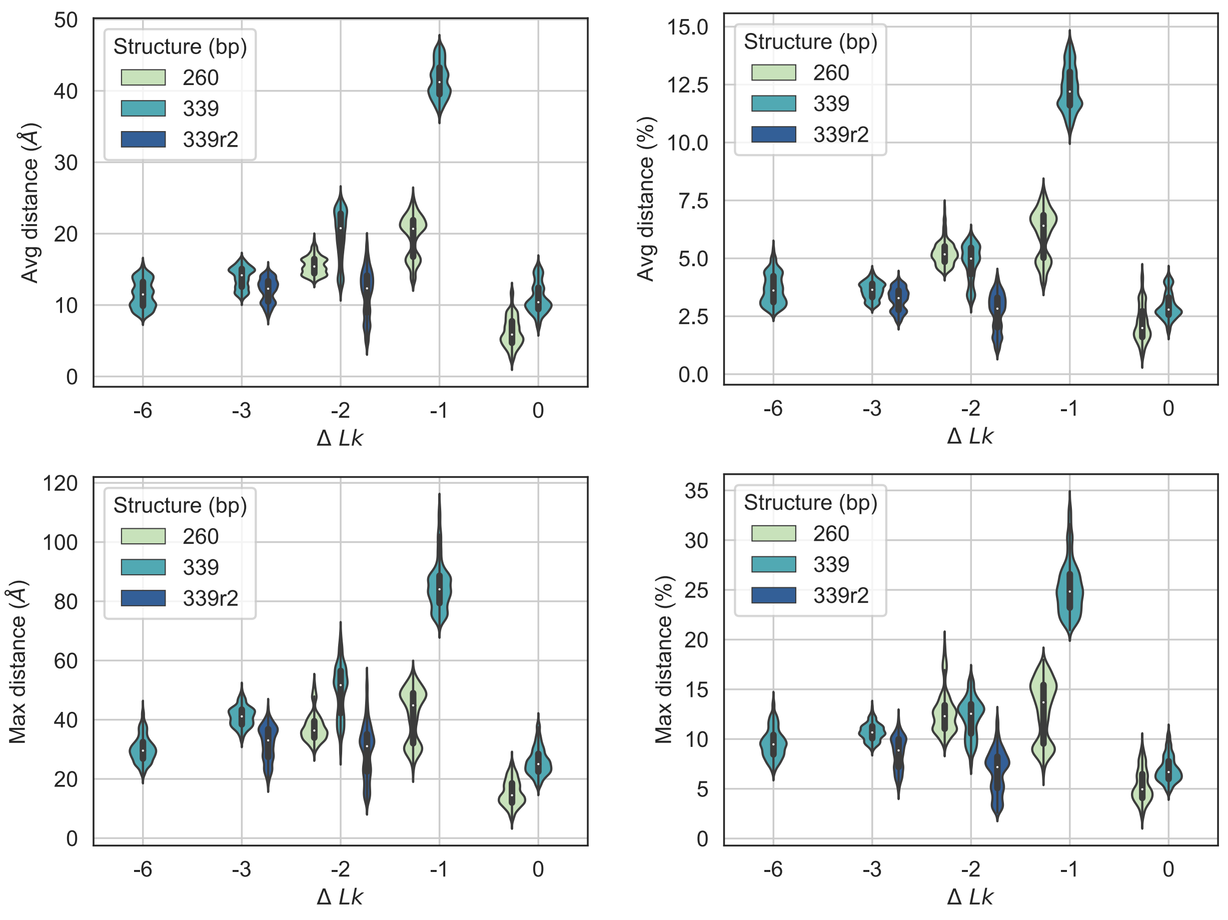


Supplementary Figure 1: In silico analysis shows a low deviation from planarity in both 260 bp and 339 bp minicircles, including for the topoisomers -2 and -3 where a second replica is run (339r2). For both systems, the average deviation from planarity at any simulation frame is less than 3 nm (~8%) for all superhelical densities except ΔLk = -1 (339 bp), where this increases to 5 nm (~15%). The deviation from planarity at any one point of the molecule is less than 20% for all molecules except ΔLk -1 and for ΔLk -1 is less than 35%.

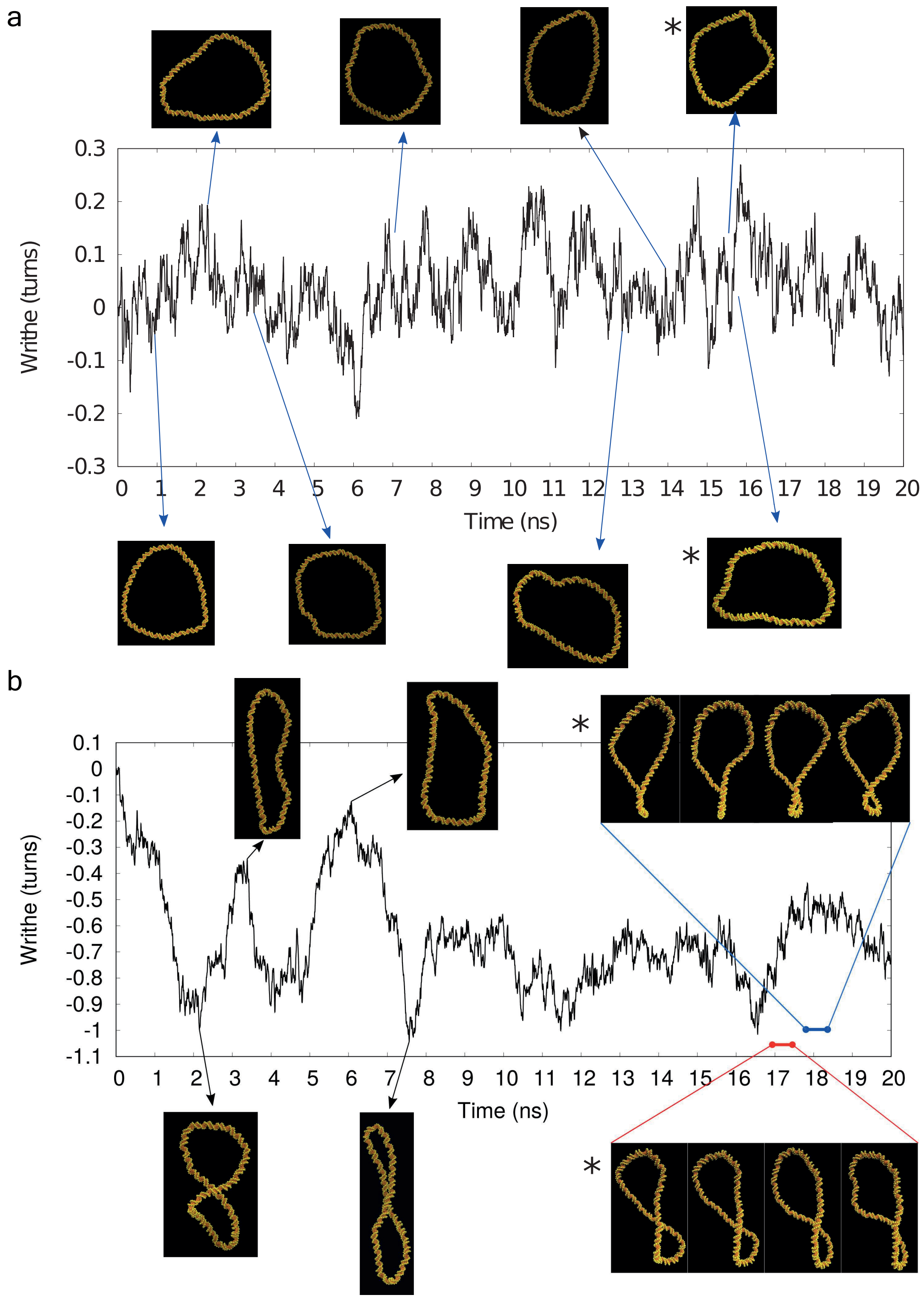


Supplementary Figure 2: Time evolution of writhe in conjunction with selected snapshots showing different molecular conformations from simulations of the relaxed topoisomer (ΔLk = 0) (a) and the negatively supercoiled topoisomer (ΔLk = -1) (b) run in implicit solvent. Asterisks mark the individual structures depicted on Fig. 1e and the chronological series selected for Fig. 1f and 1g. We observe global molecular rearrangements, specially in ΔLk = -1, and similar configurations sampled repeatedly during simulations. Remarkably, the first structure for the ΔLk = 0 minicircle resembles the average structure of the same topoisomer obtained in explicit solvent (Fig. 2a)

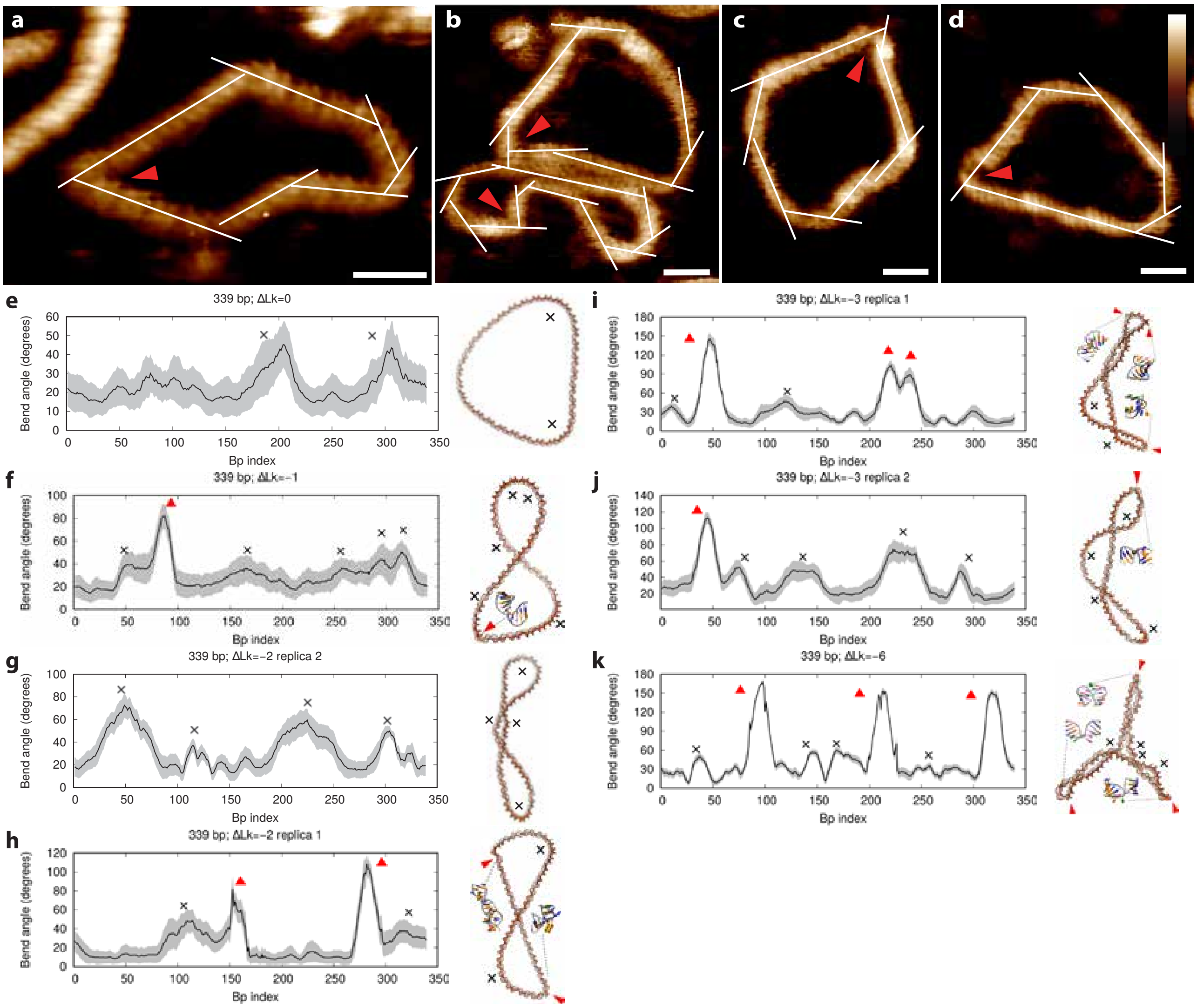


Supplementary Figure 3: Analysis of bend angles by AFM and MD simulations. a-d High-resolution AFM images of 251 (a) and 339 (b, c, d) bp minicircles, analysed using Gwyddion[1] to calculate bend angles. The bend angle for bent and kinked regions were calculated as 51 ± 14° and 106 ± 16°. Errors quoted are standard deviation (N = 33 and 5 respectively). At this resolution the minicircles show observable differences in structure, as kinked (red arrow head) or bent regions. Scale bars 10 nm, height scale (inset) 2.5 nm. e-k Bend angles for all sub-fragments 16 bp-long of MD simulations calculated by the SerraLINE program using the WrLINE profile. All peaks higher than 35$^0$ are selected for Figure 2d and are classified as B-DNA bends (black cross) or defects (red triangles) depending on whether canonical non-bonded interactions were broken or not. Grey shades correspond to standard deviations calculated along the last 30 ns of the simulations

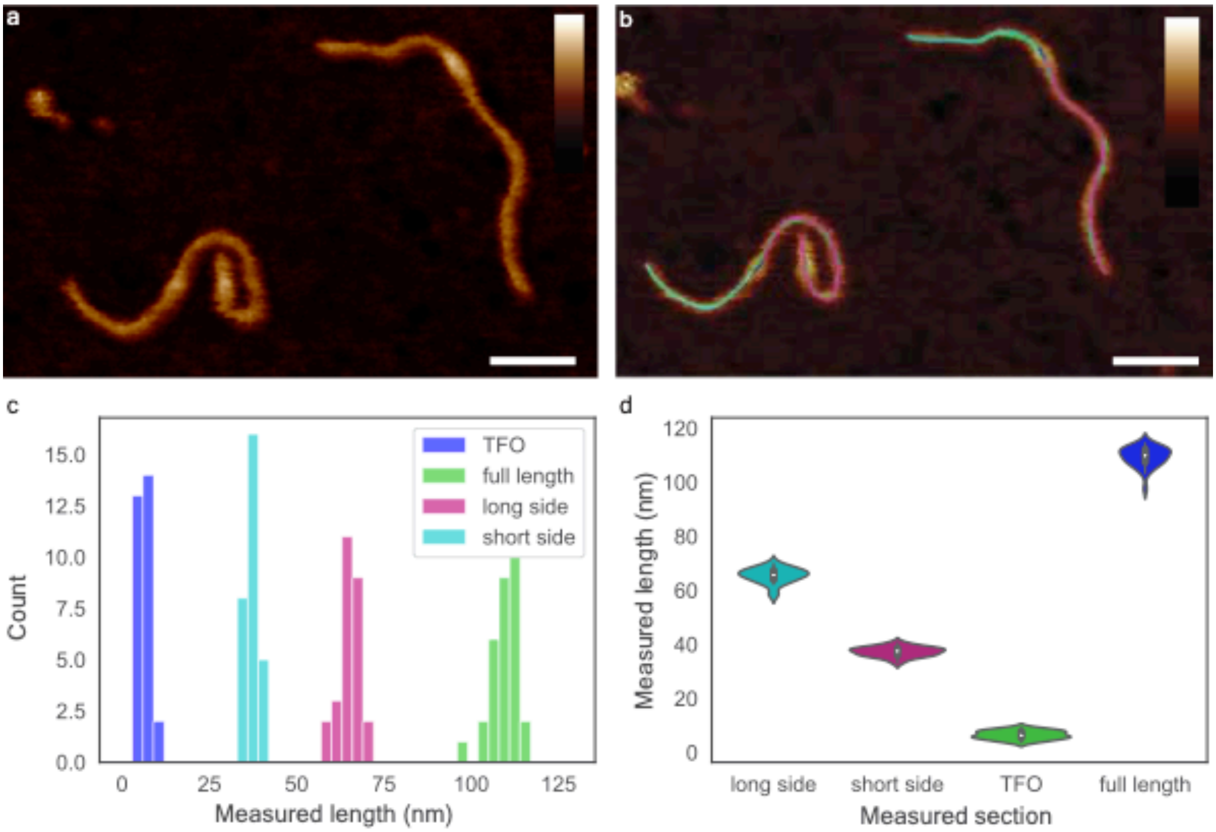


Supplementary Figure 4: Confirmation of triplex formation in 339 bp minicircles by AFM. a, AFM image of linearised 339-bp minicircles showing a protrusion at a location consistent with triplex formation. Minicircles linearised with NdeI, incubated with the triplex-forming oligonucleotide TFO1 and immobilised on mica using Poly-L-lysine in 50 mM NaOAc, pH 5.2, at room temperature. b, AFM images are hand traced in IMOD[2] to determine the length of the linearised minicircle, and the lengths of DNA either side of the protrusion. c, d, Histogram and violin plot showing the length of the TFO, full length, long and short sides. Lengths were determined by AFM to be 6 ± 2, 109 ± 4, 65 ± 3 and 37 ± 2 nm (SD) respectively for N=29 separate molecules. This is in good agreement with calculations by base-pair separation for the full length and short sides as 115 nm (339 bp) 43 nm (127 bp). Scale bars: 25 nm, height scale (scale bar inset): 3 nm (a) 4 nm (b).

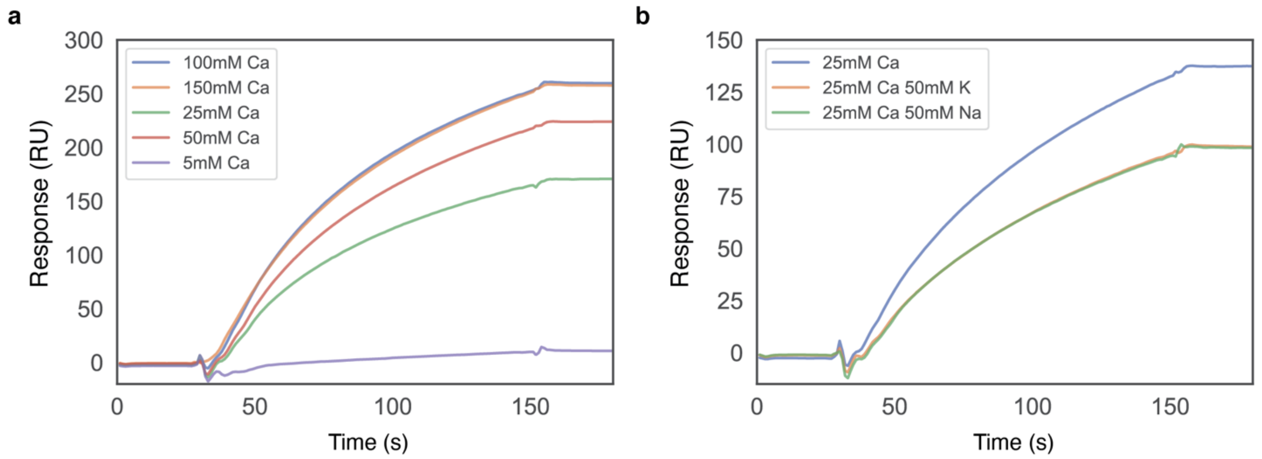


Supplementary Figure 5: Ionic concentration affects triplex formation in minicircle DNA. a, SPR shows the effect of increasing concentrations of calcium on triplex formation. The binding response increases until a maximum is observed at 100 mM. b, SPR shows the effect of monovalent cations on triplex formation. Lower binding efficiencies are observed in the presence of 50 mM sodium chloride or potassium chloride than for calcium acetate buffer alone (25 mM). In all experiments biotinylated TFO1R was immobilised to 250 RU on a streptavidin chip in a Biacore T200 instrument, and pNO1 (200 nM) was flowed over at 2 µL/min for 120 s followed by buffer only for 180 s at pH 4.6 and 25°C.

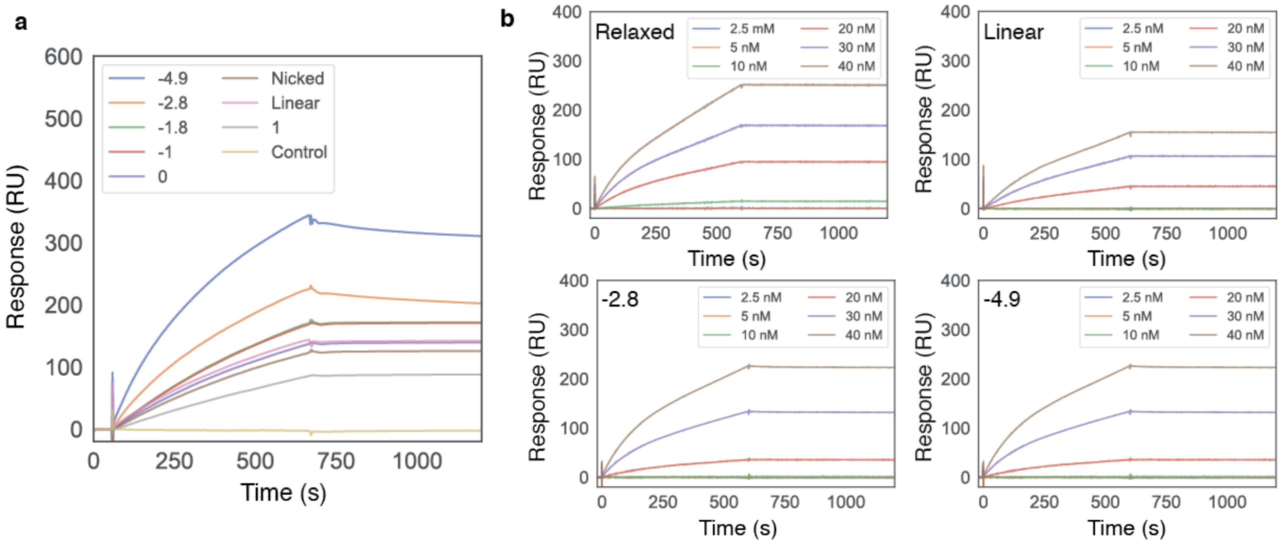


Supplementary Figure 6: Supercoiling does not substantially affect the affinity of triplex formation in minicircle DNA. a, SPR shows capture of 339 bp minicircles (ΔLk -4.9 to +1, and relaxed, nicked, linear and non-triplex forming 339 bp mincircle circle (Control) by TFO1R. b, Kinetics of the triplex binding interaction. To determine the kinetics of this interaction a range of TFOR1 concentrations (2.5, 5, 10, 20, 30 and 40 nM) were injected for each sample for 10 mins and then buffer flowed for 1 hour (for clarity, sensorgrams are only shown for 20 mins). Kinetic parameters were extracted from b, shown in Supplementary Table 1. In all experiments, biotinylated TFO1R was immobilised to 250 RU on a streptavidin chip in a Biacore T200 instrument and minicircles (50 nM) were flowed over the triplex forming sequence TFO1, immobilised on a streptavidin chip surface (2 µL/min, 600 s) in 100 mM calcium acetate, pH 4.8, 35°C.

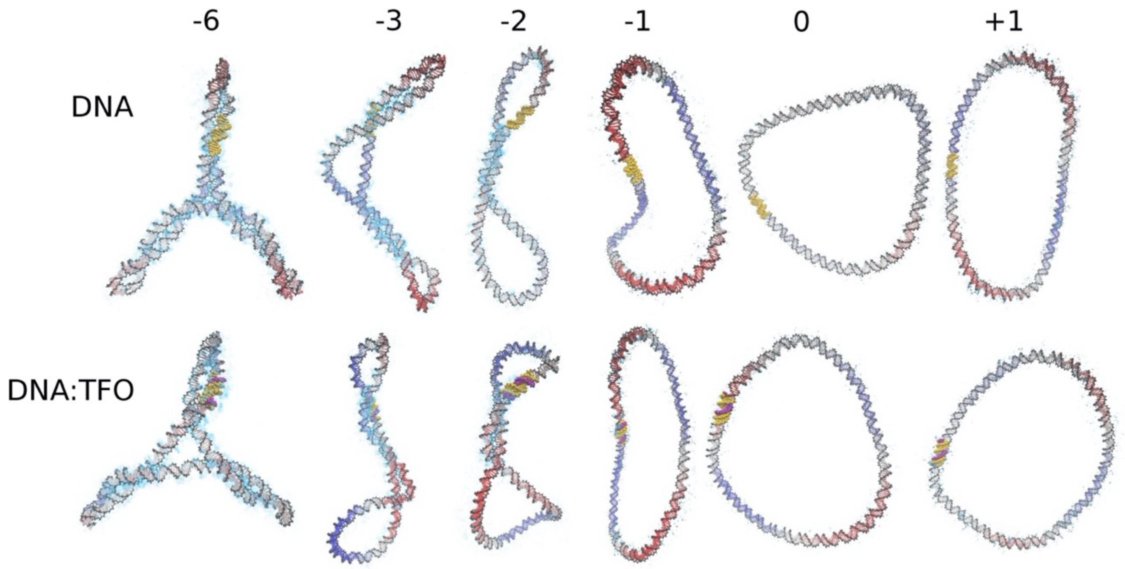


Supplementary Figure 7: Averaged structures extracted from the last 30 ns of explicitly solvated MD simulations of the 339 bp DNA mincircle topoisomers with and without the triplex-forming oligomer (TFO) together with $Ca^{2+}$-density maps (in cyan contours with an occupancy of approximately 4 times or greater the bulk concentration). Structures are colour-coded according the perspective (areas close to the reader are red, whereas those far away are in blue) so that the global shape for the writhed loop can be clearly seen. The triplex-binding site is highlighted in yellow and the TFO in purple.

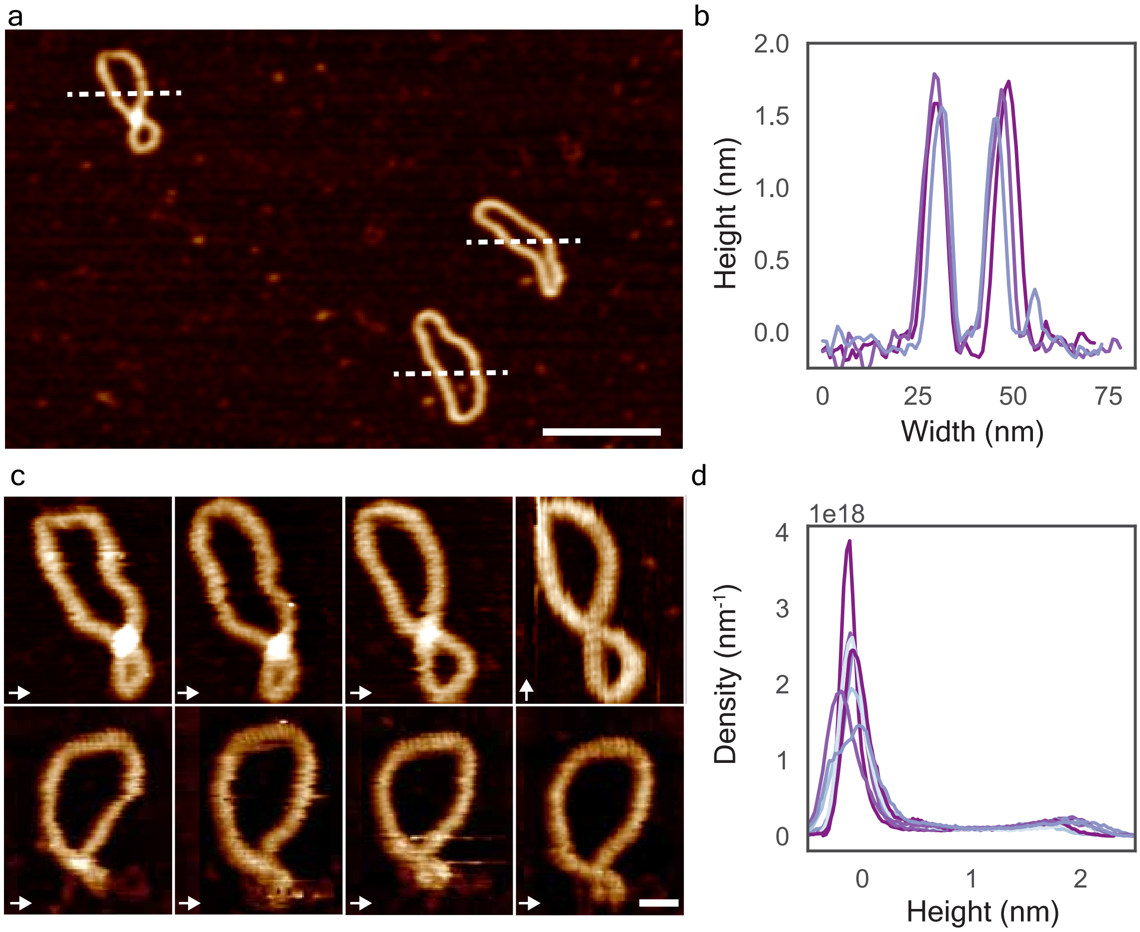


Supplementary Figure 8: Analysis of DNA minicircle compression by AFM obtained in tapping mode on a home-built microscope. a, AFM image of 339 bp DNA minicircles, dotted lines indicate location of line profile measurements. Scale bar: 50 nm, height scale 3 nm. b, line profiles taken from a, showing the mincircle height. Average heights were measured for each molecule using Gwyddion's grain measuring tool and calculated to be 1.5 ± 0.03 nm (N = 7, mean ± std). c High-resolution time-lapse imaging of the top left DNA minicircle in a, showing variation in conformation over time. d Height distributions for each image in c, showing peak heights of ~1.8 nm.

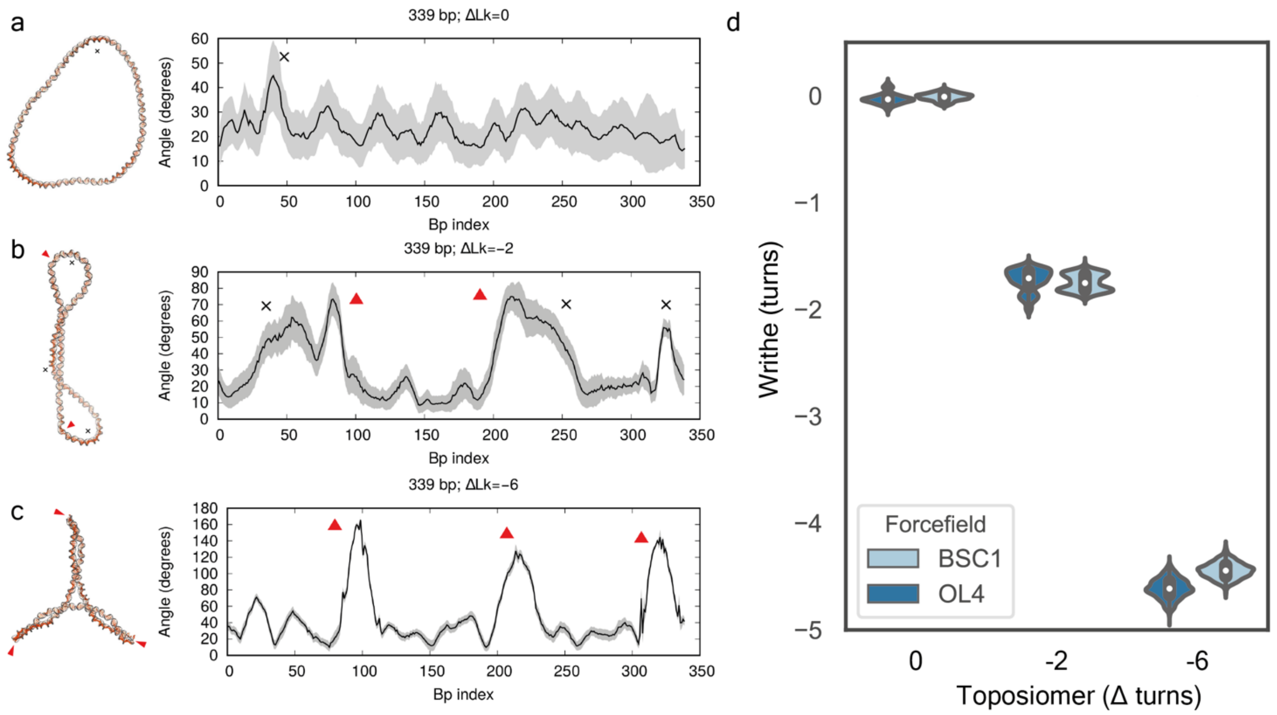


Supplementary Figure 9: OL4 and BSC1 DNA forcefields describe supercoiled DNA minicircles broadly with same structure and dynamics. a-c, Average structures and bend angles obtained from BSC1 simulations are similar to the ones obtained from OL4 simulations (see Supplementary Figure 1). Bending profiles are calculated by SerraLINE program using the WrLINE profile for all sub-fragments 16 bp-long. All peaks higher than 40$^0$ are classified as B-DNA bends (black cross) or defects (red triangles) depending on whether canonical non-bonded interactions were broken or not. Grey shading corresponds to standard deviation calculated along the last 30 ns of the simulations. d Comparison of Writhe calculations obtained from the different forcefields, OL4 and BSC1.

| 339 nr minicircle ΔLK | $k_{on}$ (ms$^{-1}$) [SE] | $k_{off}$ (s$^{-1}$) [SE] | $K_d$ (M) |
|---|---|---|---|
| -4.9 | 8.1 x $10^4$ [71] | 3.4 x $10^{-6}$ [2.1 x $10^{-8}$] | 4.2 x $10^{-11}$ |
| -2.8 | 8.3 x $10^4$ [42] | 1.9 x $10^{-6}$ [9.3 x $10^{-9}$] | 2.3 x $10^{-11}$ |
| Relaxed | 6.2 x $10^4$ [91] | 4.5 x $10^{-6}$ [2.7 x $10^{-8}$] | 7.3 x $10^{-11}$ |
| Linear | 4.8 x $10^4$ [14] | 3.6 x $10^{-7}$ [1.8 x $10^{-10}$] | 7.6 x $10^{-12}$ |
| No TFS -1.8 | NB | NB | NB |

Supplementary Table 1: Kinetics of triplex formation between TFO1R and 339 bp minicircle samples as measured by SPR. DNA analyte was flowed at 2 µL/min over a streptavidin-coated chip that had previously been loaded with TFO1R (221 RU). Running buffer was 100 mM calcium acetate (pH 4.8) and all experiments were carried out at 25 °C. The minicircle was injected at a range of concentrations from 2.5-40 nM for 600 s and the dissociation was measured for 1 h. Even with these parameters the dissociation was too slow to be accurately determined by the instrument. All data were fitted using the Biacore T200 evaluation software assuming a 1:1 binding model. The values obtained are broadly similar showing that the degree of supercoiling does not play a large effect in triplex formation when minicircles are used. (TFS = triplex-forming sequence; NB = no binding.)

| Primer name | Primer sequence |
|---|---|
| 339NEW TFONearF | 5'GAACAAACTTTCTTGTACGCGGTGGTGAGAGAGAGAGAGAGATACGACTACTATCAGCC 3' |
| 339NEW TFONearR | 5'GGCTGATAGTAGTCGTATCTCTCTCTCTCTCTCACCACCGCGTACAAGAAAGTTTGTTC 3' |
| 251 TFONearF | 5'CTTGTATACCTTTAAGAGAGAGAGAGAGAGAGACGACTCCTGCGATATC 3' |
| 251 TFONearR | 5'GATATCGCAGGAGTCGTCTCTCTCTCTCTCTCTTAAAGGTATACAAG 3' |
| TFO1R | 5'[Bt}-CTCTCTCTCTCTCTCT 3' |

Supplementary Table 2: Primers used in the formation of DNA mincircles

References:

1. Nečas, D. & Klapetek, P. Gwyddion: an open-source software for SPM data analysis. *Cent. Eur. J. Phys.* **10**, 181–188 (2011).
2. Kremer, J. R., Mastronarde, D. N. & McIntosh, J. R. Computer Visualization of Three-Dimensional Image Data Using IMOD. *J. Struct. Biol.* **116**, 71–76 (1996).
